# prolfqua: A Comprehensive R-package for Proteomics Differential Expression Analysis

**DOI:** 10.1101/2022.06.07.494524

**Authors:** Witold E. Wolski, Paolo Nanni, Jonas Grossmann, Maria d’Errico, Ralph Schlapbach, Christian Panse

## Abstract

Mass spectrometry is widely used for quantitative proteomics studies, relative protein quantification, and differential expression analysis of proteins. Nevertheless, there is a need for a flexible and easy-to-use application programming interface in R that transparently supports a variety of well principled statistical procedures. The *prolfqua* package can model simple experimental designs with a single explanatory variable and complex experiments with multiple factors and hypothesis testing. It integrates essential steps of the mass spectrometry-based differential expression analysis workflow: quality control, data normalization, protein aggregation, statistical modeling, hypothesis testing, and sample size estimation. The application programmer interface strives to be clear, predictable, discoverable, and consistent to make proteomics data analysis easy and exciting. Furthermore, the package implements benchmark functionality that can help to compare data acquisition, data preprocessing, or data modeling methods using a gold standard dataset. Finally, we show that the implemented methods allow sensitive and specific differential expression analysis. The *prolfqua* R package is available on GitHub https://github.com/fgcz/prolfqua, distributed under the MIT licence, and runs on all platforms supported by the R free software environment for statistical computing and graphics.

## 1 Introduction

> To paraphrase provocatively, ‘machine learning is statistics minus any checking of models and assumptions’.
>
> – Brian D. Ripley, useR! 2004, Vienna.

Proteins carry out most crucial functions and give structure to living cells. Hence, measuring changes in protein abundance is the subject of active research (Vidova and Spacil 2017). Bottom-up mass spectrometric methods, where proteins are specifically and reproducibly digested into protein fragments - peptides, are employed to identify and quantify proteins in complex samples containing hundreds to thousand of proteins (Bubis et al. 2017; Veiga Leprevost et al. 2020). The peptides are first separated by their chemical and physical properties using liquid chromatography (LC). Afterward, they are ionised, weighed, identified, and quantified using mass spectrometric techniques, e.g., electro-spray-ionization mass spectrometry (ESI-MS). Finally, peptide identification is achieved by fragmenting and matching the measured fragment masses to theoretical masses computed from known peptide sequences (Eng et al. 2015; Yu, Li, and Yu 2016; Kong et al. 2017). For quantification, intact peptide ions (Yu et al. 2020; Cox and Mann 2008) or products of peptide ion fragmentation (Röst et al. 2014; Demichev et al. 2020) are counted and aggregated to obtain peptide abundances. Finally, the identified and quantified peptides are assigned to proteins based on protein sequence information.

Proteomics quantification experiments enable monitoring relative abundances of thousands of proteins in biological samples. Most studies use parallel-group designs, where one or many treatment groups are compared to the control group (Leeuw et al. 2022; Laubscher et al. 2021). More recently, more complex experimental designs with an increasing number of samples are studied, e.g., factorial designs and longitudinal studies (time series), sometimes with repeated measurements on the same subject (Tan et al. 2022; Meier-Abt et al. 2021). The data can be modeled using linear fixed-, mixed-, or random-effects models (Bates et al. 2015). Based on these models, tests can be applied to examine whether specific factors and factor interactions are significant, e.g., it can be tested if differences in protein abundance between groups are statistically significant.

An important aspect of mass spectrometric data are missing peptide and protein quantifications. Rubin (1976) classified missing data problems into three categories: missing completely at random (MCAR), missing at random (MAR), and missing not at random (MNAR). For instance, in data-dependent acquisition (DDA) MS, only a limited number of MS1 signals are selected for fragmentation and identified. Peptide quantification algorithms transfer identification information between MS1 features in different samples by aligning the data using retention time and mass information, thus reducing the amount of missing data. Nevertheless, highly abundant proteins can suppress the detection of other proteins in a sample, a MAR mechanism. Furthermore, a weak correlation between the number of missing measurements in a group and the abundance of the quantified protein can be observed, which is caused by the limit of detection (LOD), a MNAR mechanism (McGurk et al. 2020).

Several data analysis packages exist to model mass spectrometry protein quantification experiments, e.g., *limma* (Ritchie et al. 2015), *MSstats* (Choi et al. 2014), *PECA* (Suomi and Elo 2017), *msqrob2* (Goeminne, Gevaert, and Clement 2016) or *proDA* (Ahlmann-Eltze and Anders 2020), to mention some, all implemented in R. Each of them has some unique features, for example, *MSstats* infers the model and generates the model formula from the sample annotation, allowing users with limited statistical knowledge to perform differential expression analysis (DEA). At the same time, *limma* allows to specify a design matrix using a linear model formula and implements the empirical Bayes variance shrinkage method. The package *PECA*, performs a roll-up of peptide level differences and peptide level *p*-value estimates, obtained from *limma* or *PECA*, to protein level estimates. Furthermore, *msqrob2* implements two models: robust linear models fitted to protein intensities and robust ridge regression fitted to peptide intensities, and combines them with empirical Bayes variance shrinkage. The *proDA* package implements a linear probabilistic dropout model to account for missing data without imputation. *MSstats* handles missing data by feature filtering and using imputation. Other means of modelling missing observations are the Hurdle model discussed by Goeminne et al. (2020), while the R package *proDA* models missingness using probabilistic dropout models (Ahlmann-Eltze and Anders 2020).

When analyzing parallel-group designs using a single explanatory variable, and contrasting groups, we can use all R packages; but, we can use only some of them if we want to model factorial designs or repeated measurements. Table 1 gives an overview of the models and features supported by these packages. We see that packages such as *limma* and *proDA* allow us to fit a comprehensive variety of models and test various hypotheses; however, in-depth knowledge of design matrix specification using the R formula interface is required (Law et al. 2020).

**Table 1:**
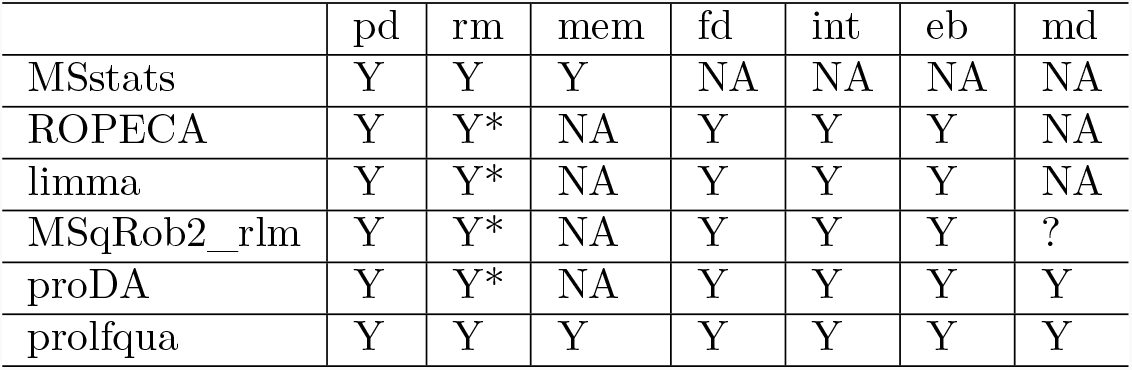
Rows - R package, Columns - types of models supported: pd - parallel design, rm - repeated measurements, fd - factorial design, int - interactions among factors, mem - mixed effect models, eb - empirical Bayes, md - missing data modelling. Y - yes, * - repeated measurements are modeled using a fixed effect. ? - the hurdle model was published but is not available in the msqrob2 package.

When developing the R package *prolfqua* we were inspired by the R package *caret* (Kuhn 2008), which enables us to call various machine learning (ML) methods, and makes selecting the best ML algorithm for your problem easy. We aimed for a similar R package for the differential expression analysis of proteomics data. However, after examining the R packages for modeling proteomics quantification data, we observed that model specification, input, and output formats differ widely. At the same time, they have in common that: they fit linear models either to peptide or protein intensities, determine differences among groups, and afterward apply empirical Bayes variance shrinkage. Therefore, the revised goal was to provide a modular object-oriented (OO) design, with R linear models as a core, where we can add features such as *p*-value aggregation, e.g., *PECA*, variance shrinkage, or modeling of missing observations.

Furthermore, the functionality of *prolfqua* also includes methods specific to proteomics data. For example, we implemented strategies to estimate protein intensities from peptide intensities: top N (Grossmann et al. 2010), Tukey’s median polish (Tukey and others 1977), robust linear models (Goeminne et al. 2020). Furthermore, peptide or protein abundances can be scaled and transformed, using robust scaling, *quantile* normalization or *vsn* to remove systematic differences among samples and heteroscedasticity. In this respect, *prolfqua* can be compared with other R packages such as *DEP* (Zhang et al. 2018) or *POMA* (Castellano-Escuder, Andrés-Lacueva, and Sánchez-Pla 2021) which support the entire differential expression analysis pipeline.

We also implemented functionality and use the Ionstar (Shen et al. 2018) dataset to benchmark the modeling methods implemented within *prolfqua* and to compare our results with those of *MSstats* and *proDA*. Since group sizes are relatively small, typically with four or five subjects per group, the high power of the tests is a relevant criterion to assess the modeling method. The quantified proteins can be ranked using the estimated fold-change, *t*-statistics, or scaled *p*-value, and afterward subjected to gene set enrichment (GSEA) or over-representation analysis (Subramanian et al. 2005) to determine up or down-regulated groups of proteins. Furthermore, the statistical model must provide an unbiased estimate of the false discovery rate (FDR) to manage expectations when selecting protein lists for follow-up experiments. We will use the partial area under the receiver operator curve (ROC) to assess the power of the tests and compare the FDR with the false discovery proportion (FDP).

## 2 Methods

### 2.1 Implementation

We store all the data needed for analysis in a tidy table, i.e., every column is a variable, every row is an observation, and every cell is a single value (Wickham 2014). Using an R6 (Chang 2020) configuration object (Figure 1), we specify what variable is in which column, making it easy to integrate new inputs in *prolfqua* if provided as a tidy table. For example, to visualize tidy Spectronaut(Bruderer et al. 2015), DiaNN(Demichev et al. 2020), or Skyline(MacLean et al. 2010) outputs, or data in *MSstats*(Choi et al. 2014) format, only a few lines of code are needed to update the *prolfqua* `AnalysisTableConfiguration` configuration. The configuration encapsulates the differences in column names among the various input formats, and enables the use of the methods without additional arguments. We show an example code for creating an MSFragger(Yu et al. 2020) configuration in the Appendix. For popular software like MaxQuant(Cox and Mann 2008), or MSFragger, which stores the same variable, e.g., intensity, in multiple columns, one for each sample, we implemented methods that transform the data into tidy tables. Relying on the tidy data table enables us to interface with many data manipulation, visualization, and modeling methods, implemented in base R (R Core Team 2021) and the tidyverse (Wickham et al. 2019), easily. We use R6 classes to structure the functionality of the package (see Figure 1 and Figure 2). R6 classes are well supported by command-line completion features in RStudio (RStudio Team 2022), and help to implement argument free functions (Figure 6.

**Figure 1:**
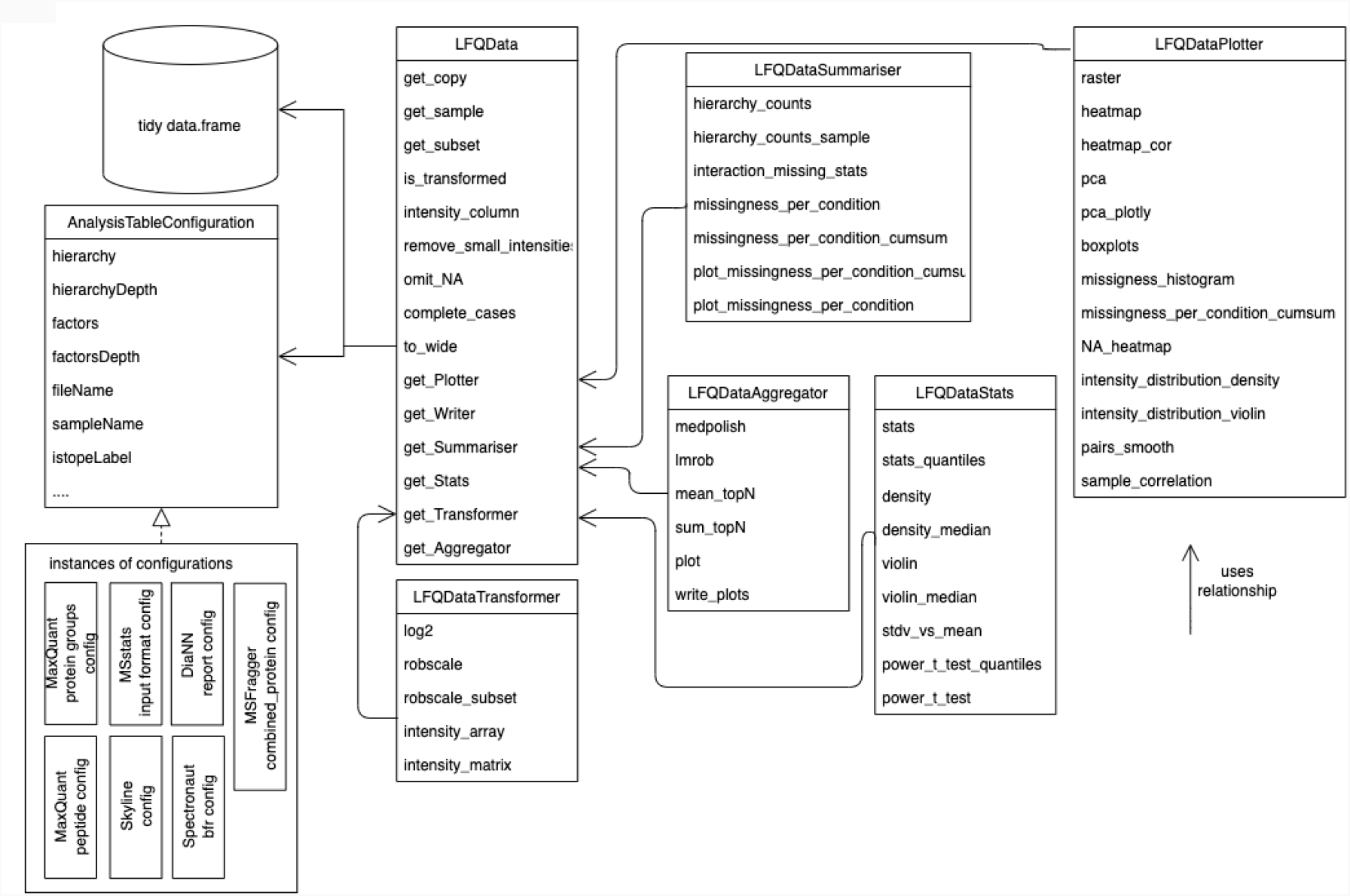
Class Diagram of classes representing the proteomics data. The LFQData class encapsulates the quantitative proteomics data stored in a tidy table. An instance of the AnalysisTableConfiguration class specifies a mapping of columns in the tidy table. The LFQDataPlotter class and other classes decorate the LFQData class with additional functionality. Similarly, the `LFQDataStats` and `LFQDataSummary` reference the `LFQData` class and group methods for variance and sample size estimation or summarizing peptide and protein counts.

**Figure 2:**
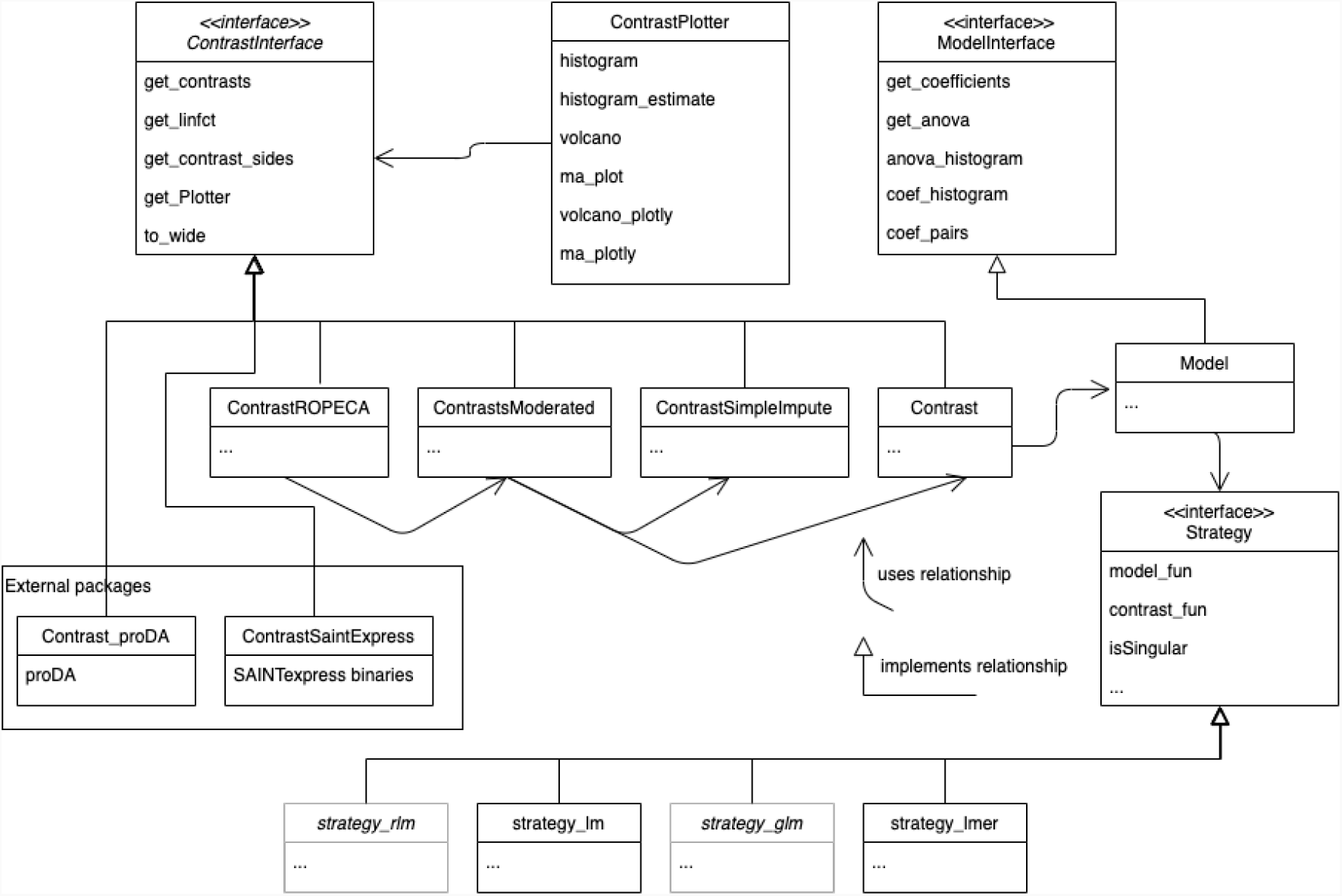
The UML (unified modeling language) diagram of modeling and contrast related classes. Different strategies, e.g., *lm, lmer*, and *glm* (Table 3), can fit various types of models. The model builder method fits the statistical model given the data and a strategy. The obtained model can then be used for the analysis of variance or to estimate contrasts. All classes estimating contrasts implement the *Contrasts* interface. Results of external tools, e.g., SAINTexpress, can be adapted by implementing the Contrasts Interface.

In addition, we implement features specific to high throughput experiments, such as the experimental Bayes variance and *p*-value moderation, which utilizes the parallel structure of the protein measurements and the analysis (Ritchie et al. 2015). We also compute probabilities of differential protein regulation based on peptide level models (Suomi and Elo 2017). We used R6 classes to encapsulate the statistical modelling functionality in *prolfqua* (see Figure 2). We specify a contrast interface `(ContrastsInterface)`. Several implementations allow to perform differential expression analysis given linear or mixed effect models `(Contrasts),` variance shrinkage `(ContrastsModerated),` or to impute contrasts in cases when observations are missing for an entire group of samples `(ContrastsSimpleImpute)`. Further implementations of the interface encapsulate and integrate differential expression analysis results of external tools such as *proDA* or of SAINTexpress (Teo et al. 2014) used to analyze data from protein interaction studies.

### 2.2 Dataset for benchmarking

To evaluate the performance of differential expression analysis, we use the IonStar benchmark dataset(Shen et al. 2018), available from the Proteomics Identifications Database (PRIDE) identifier PXD003881. *DH*5*α E. coli* lysate was spiked in human pancreatic cancer cells (Panc-1) lysate at five different levels: 3% *E. coli*, 4.5% *E. coli*, 6% *E. coli*, 7.5% *E. coli* and 9% *E. coli*. We annotated these dilutions from smallest to largest with the letters *a, b, c, d, e*. By comparing the various dilutions, we obtain different effect sizes, e.g., when comparing dilution *e* (9%) against dilution *d* (7.5%), the expected difference is 1.2 for E. coli proteins and 1 for human proteins. There are four technical replicates for each dilution, and hence 20 raw files in total. We processed the raw data using MaxQuant (Cox and Mann 2008) version Version 1.6.10.43, with MaxQuant default settings. The text files generated by MaxQuant are available in the *prolfquadata* R package (Wolski 2021). To compare the performance of the various methods implemented in *prolfqua* we use only the contrasts resulting in minor differences Δ = (1.20, 1.25, 1.30, 1.50), because for bigger differences all models perform similarly.

### 2.3 Data preprocessing for model comparison

The peptide abundances (from the MaxQuant *peptide*.*txt* file) were *log*_2_ transformed and subsequently scaled, where median and the mean average deviation (mad) were obtained from the human proteins only. We removed *one hit wonders*, i.e., proteins with a single peptide assignment. Protein abundances are inferred from the peptide intensities using Tukey’s median polish. Finally, we fitted the fixed effect linear models, the dropout model *proDA* to protein abundances, the mixed effect linear model, and the *PECA* model to peptide intensities.

### 2.4 Benchmark metrics

The IonStar dataset contains *H. sapiens* proteins with constant concentrations and *E. coli* proteins with varying concentrations. We know that for *H. sapiens* proteins, the difference *β* between two dilutions should be *β* = 0, while for *E. coli* proteins, we know that the difference between dilutions should be *β ≠* 0.

We can use various statistics to examine the alternative hypothesis *β* ≠ 0: the contrast estimate, i.e. the log_2_ fold-change *β*, the *t*-statistic 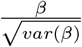, or the *p*-value and moderated *p*-value. For each statistic and each value of the statistics we then compute a confusion matrix (see Table 2). From the confusion matrix we obtain measures such as true positive rate (*TPR*), false positive rate (*FPR*), or false discovery proportion (*FDP*) which are given by:
with

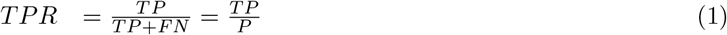

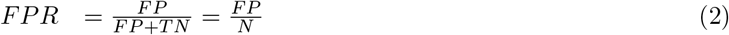

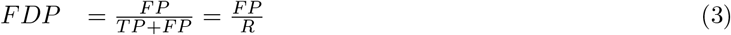

By plotting the *TPR* versus the *FPR* we obtain the receiver operator characteristic curve (ROC curve). The area under the curve (AUC) or partial areas under the curve (pAUC), at various values of the *FPR*, are measures of performance derived from the ROC curve. By using these measures we can compare the performances of the statistics produced by the various methods examined.

**Table 2:** Confusion matrix, TP - true positive, FP - false positive, FN - false negative, TN - true negatives, P - all positive cases (all *E. coli* proteins), N - all negative cases (all *H. sapiens* proteins), m - all proteins.

| Prediction \ Truth | E.coli | H.sapiens | Total |
| --- | --- | --- | --- |
| beta != 0 | TP | FP | R |
| beta == 0 | FN | TN |  |
| Total | P | N | m |

In order to compute the confusion matrices based on the *p*-value we first need to rescale it (see Equation (11)). Thus, the *p*-value is a function of the *t*-statistics and the degrees of freedom.

### 2.5 Modelling

#### 2.5.1 Robust scaling of the data

Välikangas, Suomi, and Elo (2016) discuss and benchmark various methods of peptide or protein intensity normalization, such as variance stabilizing normalization (Huber et al. 2002) or quantile normalization (Bolstad et al. 2003). In this work, we use a robust version of the z-score, where instead of the mean 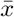 we use the median, and instead of the standard deviation 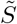 we use the median absolute deviation (mad):

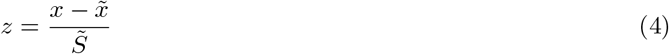

Because we need to estimate the differences among groups on the original scale, we must multiply the *z*-score by the average variance of all the *n* samples in the experiment.

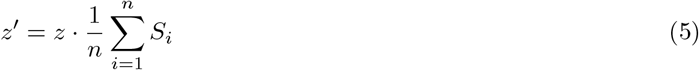

To apply this transformation, we need to estimate two parameters per sample, therefore it works for experiments with thousands of proteins and experiments where only a few hundred proteins per sample are measured. For the Ionstar dataset, we used the intensities of *H. sapiens* proteins, whose concentrations do not change, to determine 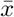 and *S* and then applied it to all the intensities (including *E. coli*) in the sample.

#### 2.5.2 Estimating differences between groups

Given a linear model *y* = *βX*, we can compute the difference *β*_*c*_ between two groups by a linear combination *c* of linear model parameters *β*, where *c* is a column vector with as many elements as there are coefficients *β* in the linear model. If *c* has 0 for one or more of its rows, then the corresponding coefficient in *β* is not involved in determining the contrast (Irizarry and Love 2018).

The difference estimate *β*_*c*_, is given by the dot product:

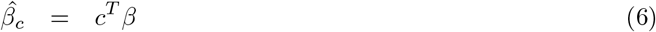

and the variance of *β*_*c*_ by:

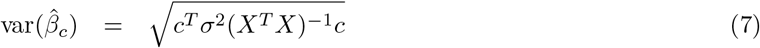

with *X* being the design matrix. The degrees of freedom for the contrast are equal to the residual degrees of freedom of the linear model. For estimating contrasts from mixed effects models we used the function contest implemented in the R package *lmerTest* and used the Satterthwaite (Kuznetsova, Brockhoff, and Christensen 2017) method to estimate the denominator degrees of freedom. These methods are available in the class `Contrast` (see Figure 2)

#### 2.5.3 Determining linear parameter combinations for treatment comparison

The package *prolfqua* provides a function to determine the vector of *parameter* weights *c*, from a linear models and a contrast specification string.

The linear model below explain the observed protein abundances using the explanatory variables `factor_1` and `factor_2` and the interaction among them `factor_1:factor_2,`

~~~
## Intensity ∼ factor_1 + factor_2 + factor_1:factor_2
~~~

then the contrasts among `group_1` and `group_2` defined by factor_1 can be specified using the string below.

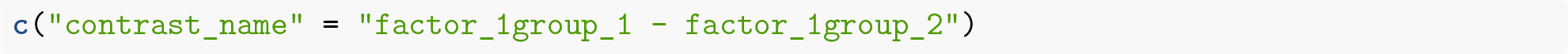

Furthermore, contrasts for subgroups can be specified using the code below,

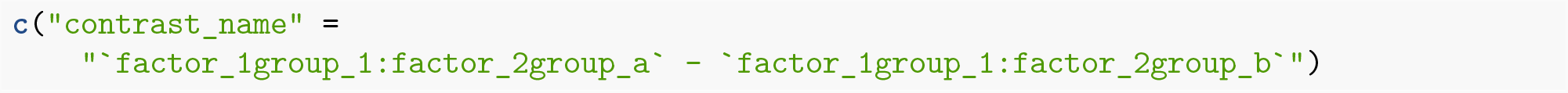

where `factor_x` is the name of the explanatory variable, and `group_x` are group labels.

The following code shows an example where we specify two contrast: the first to compare Cells of type CMP/MEP with cells of type HSC, and the second to compare therapy NO with therapy NU for the celltype CMP/MEP (Meier-Abt et al. 2021). Finally, we compute the array of weights defining the contrast, used to multiply the model coefficient *β*.

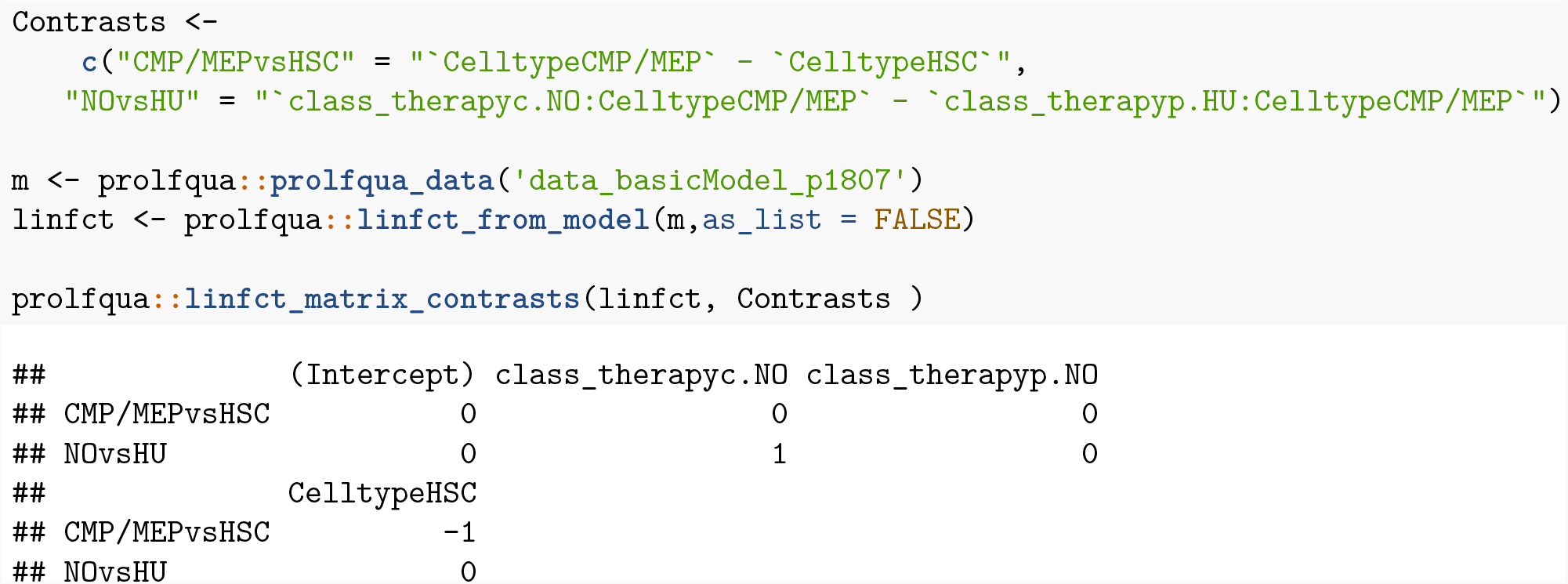

#### 2.5.4 Contrast estimation in presence of missing data

Missing observations lead to different group sizes, which results in unbalanced designs. Linear and mixed effect models can handle unbalanced designs, as long as one observation per group is available, they will produce unbiased estimates, and no imputation is needed.

If there is no observation in a group, we assume that the protein is unobserved because the protein abundance is below the limit of detection (LOD) of the MS instrument. In this case, we will impute the mean using the protein abundance at the detection limit *A*_*LOD*_. We estimate the abundance at the detection limit using the abundances of the proteins observed only in one sample of a group of samples. Then, we compute the median of the distribution and use it as the group mean if a protein is absent in a treatment group.

When computing differences Δ among two groups *a* and *b*, we will use the group mean *ā* or 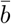 estimated from the data, or if no observations are present in a group, we use *A*_*LOD*_, e.g., If there are no observations in group *b* then :

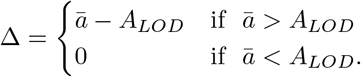

To estimate the variance, we assume that the variance of the protein is constant in all the groups and use the pooled variance based on all data:

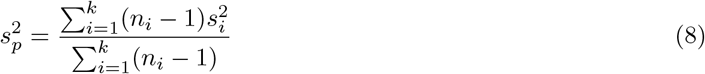

with *n*_*i*_ the number of observations, and *s*_*i*_ the standard deviation in each group. The standard deviation for the *t*-statistics is then given by:

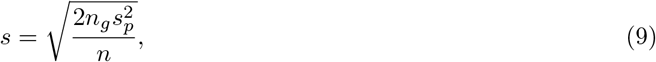

Where, *n*_*g*_ is the number of groups and *n* is the number of observations. If variance can not be estimated for a protein, we will use the median pooled variance of all the proteins.

This methods are implemented in the class `ContrastSimpleImpute` (see Figure 2).

#### 2.5.5 *p*-value moderation

From the linear and the mixed effect models, we can obtain the residual standard deviation *σ*, and degrees of freedom *df*. Smyth (2004) discuss how, to use the *σ* and *df* of all models to estimate a prior *σ* and prior *df*, and posterior 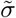. These can be used to moderate the *t*-statistics by:

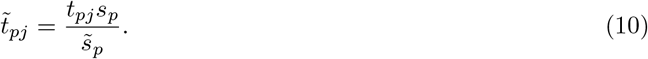

We implemented this method in the class `ContrastModerated` (Figure 2).

#### 2.5.6 Summarizing peptide level differences and p-values on protein level

To summarize peptide level models to protein models, we apply the method suggested by Suomi and Elo (2017) that use the median scaled *p*-value of the peptide models and the cumulative distribution function of the Beta distribution (CDF) to determine a regulation probability of the protein.

To obtain the 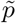 of a protein, we first rescaled the peptide *p*-values by taking the sign of the fold-change 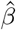 into account, i.e.:

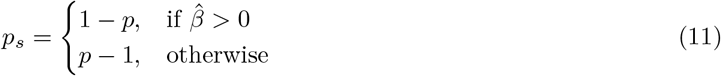

Afterwards, the median scaled *p*-value 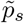 is determined and using the transformation below, transformed back onto the original scale:

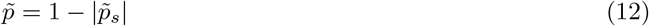

Because we use the median, with the i-th order statistic 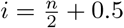, we parametrize the CDF of the Beta distribution with 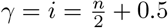 and 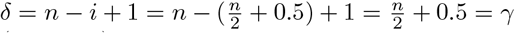. We implemented this method in the class `ContrastROPECA` (Figure 2).

## 3 Results and Discussion

### 3.1 Example analysis workflow

The code snippets in this section demonstrate how a differential expression analysis workflow can be implemented using the *prolfqua* R package. To speed up the computation of these examples, we use a random subset of the Ionstar dataset containing 163 proteins and 1258 peptides. Peptide abundances are log_2_ transformed and robust z-score scaled using the method `robscale.` Using the `LFQDataPlotter` class, we show the distribution of the normalized peptide abundances in Figure 3 Panel A. Afterwards, protein intensities are estimated from peptide intensities using Tukey’s median polish. Figure 3 Panel B shows the peptide intensities and the estimated protein intensities. Next, we compute the standard deviation of all the proteins in each group and display their distribution using violin plots (Panel C). Finally, we create a boxplot (Panel D) showing the abundance of one protein.

**Figure 3:**
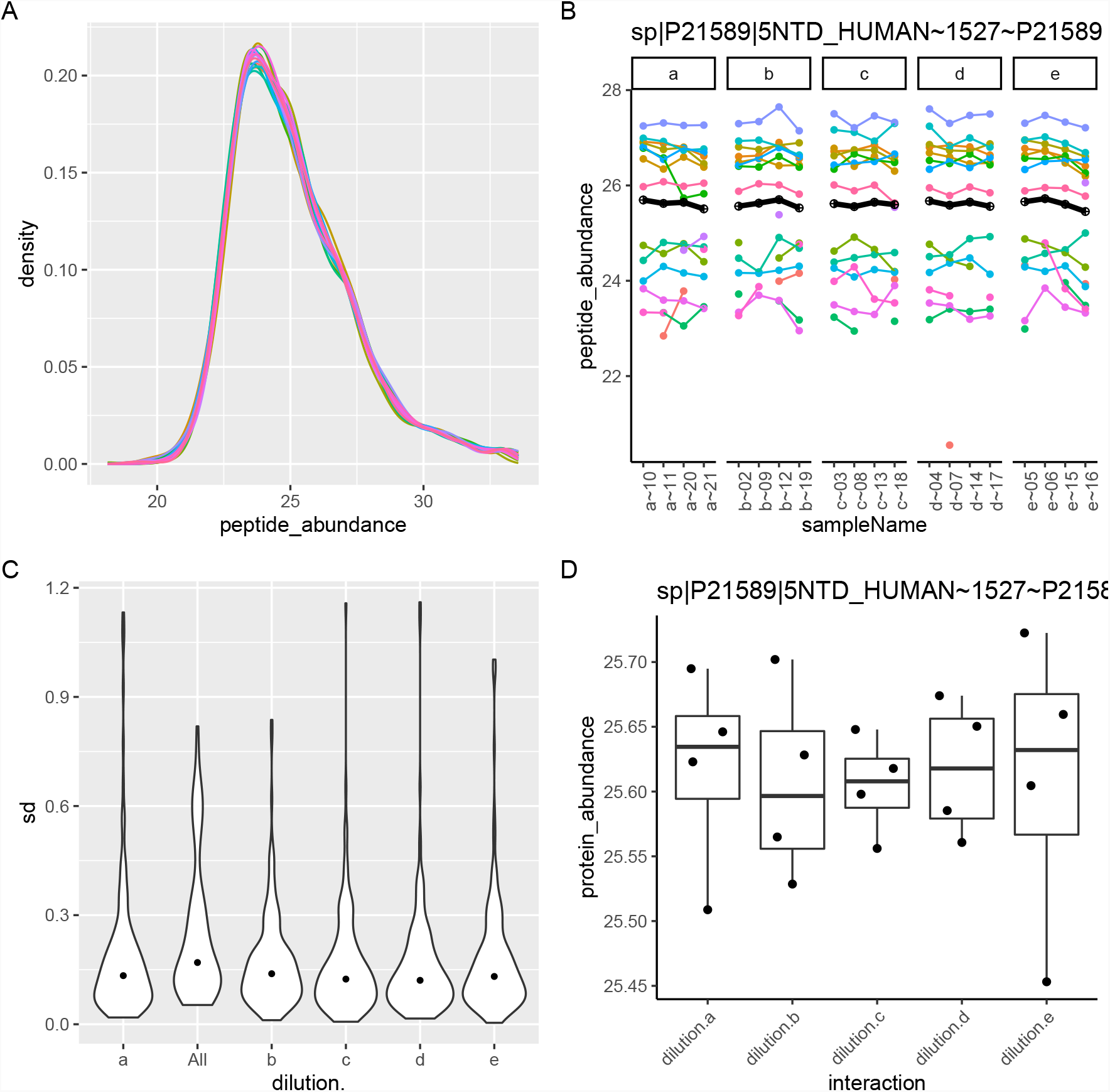
Panel A - Peptide intensity distributions for 20 samples. For each sample a line with a different colour is shown. Panel B - Peptide intensities for protein *5NTD* are shown using a line of different colour, and the protein intensity estimate is shown using a fat black line, Panel C - distribution of standard deviations of all proteins in each dilution group (*a, b, c, d, e*) and overall (all), Panel D - Distribution of protein intensities for protein *5NTD*.

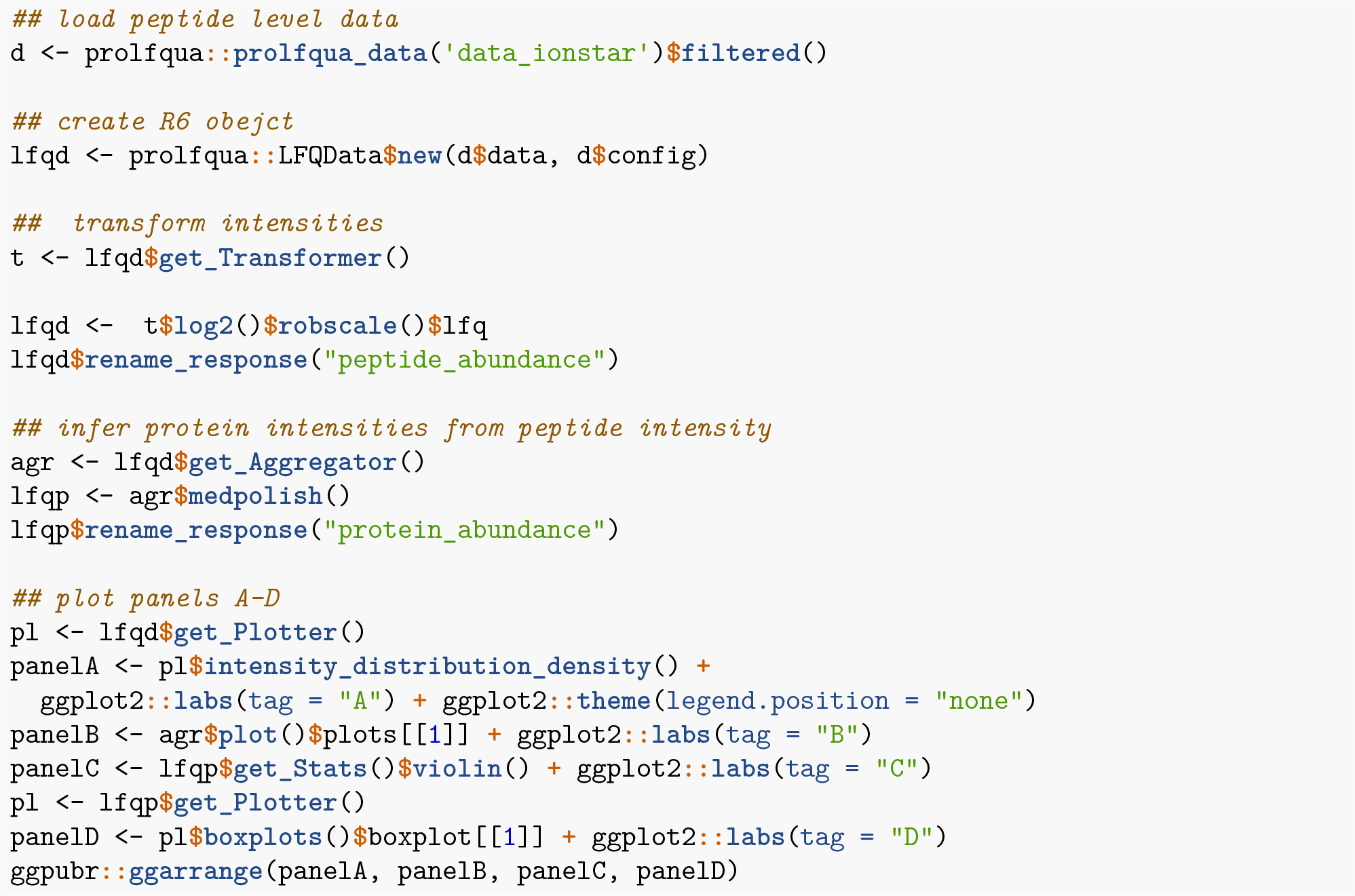

The following code example illustrates how we compute differences among groups. First, the linear model and the differences are specified. Afterward, the model is fitted to the data using the `build_model` function. Next, we estimate the contrasts either from the linear model using the Contrasts class or directly from the data using the `ContrastsSimpleImpute` class. Afterward, we apply *t*-statistic moderation using the `ContrastModerated` class. Finally, the `addContrastResults` function merges both sets of contrast estimates, preferring the one obtained from the linear model if both are available. Then we create the plots shown in Figure 4. Panel A shows the distribution of the *p*-values, while Panel B shows the volcano plot for each comparison.

**Figure 4:**
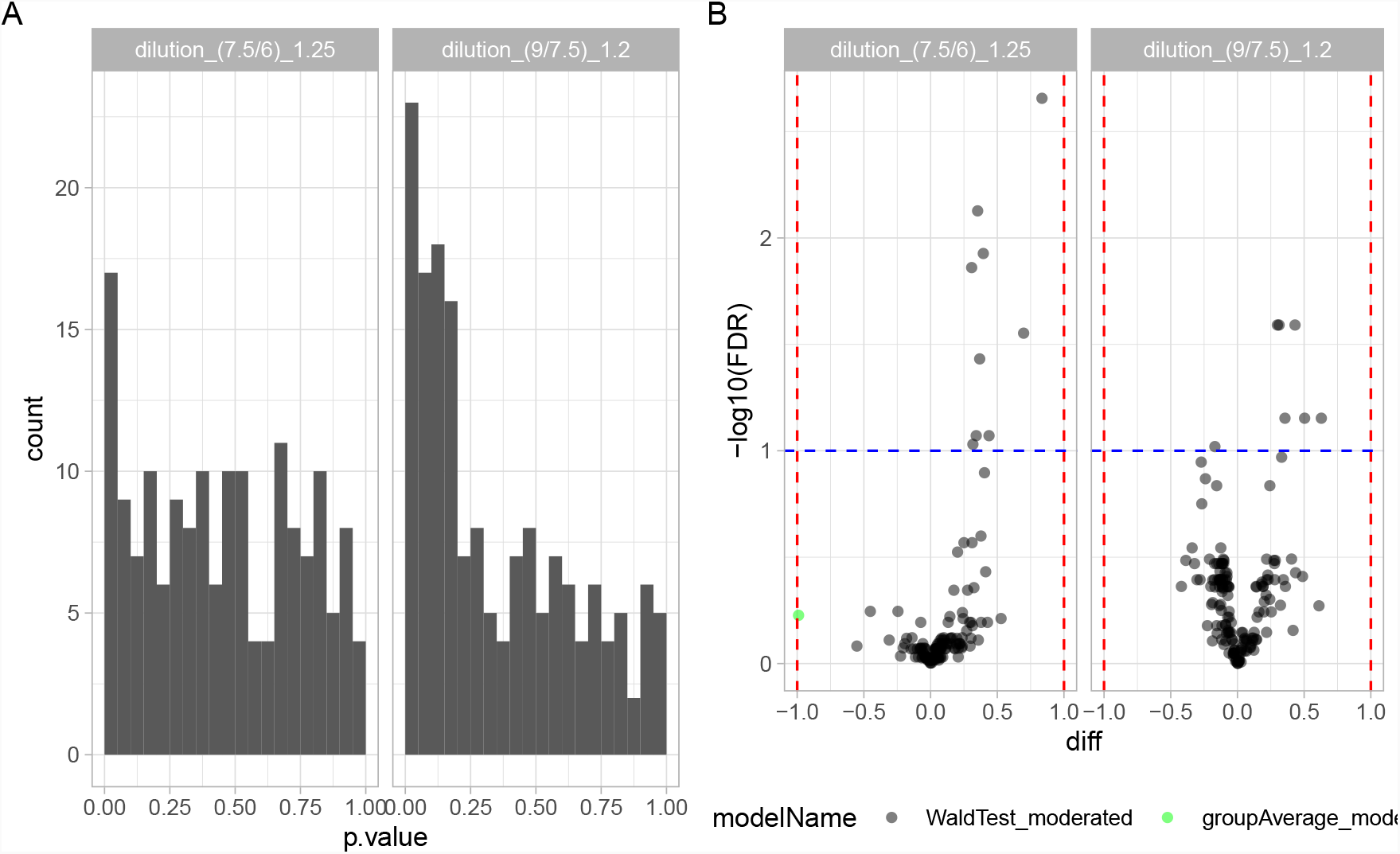
Panel A - Histogram showing the distribution of p-values for 163 proteins. Panel B - Volcano plot showing *−* log_10_ transformed FDR as function of the difference between groups for 163 proteins.

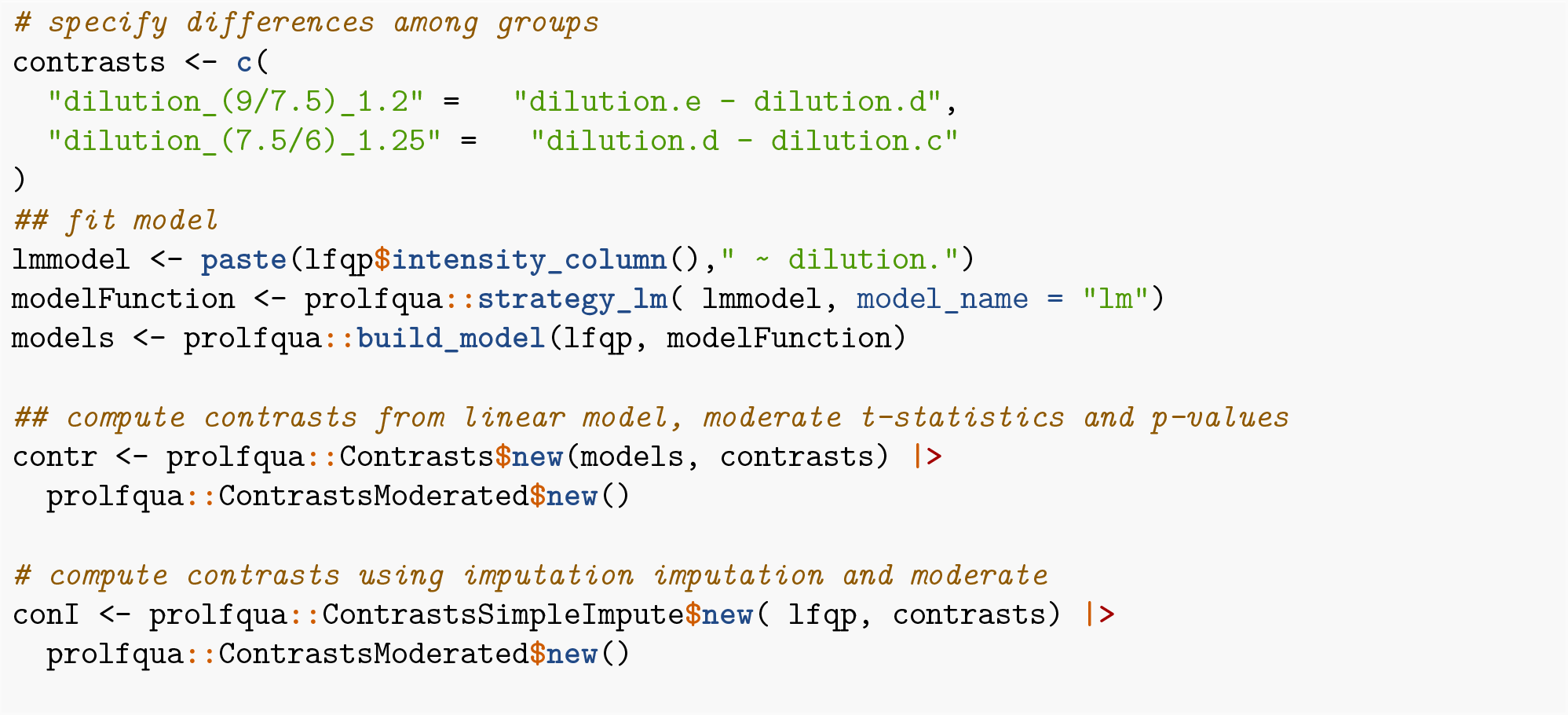

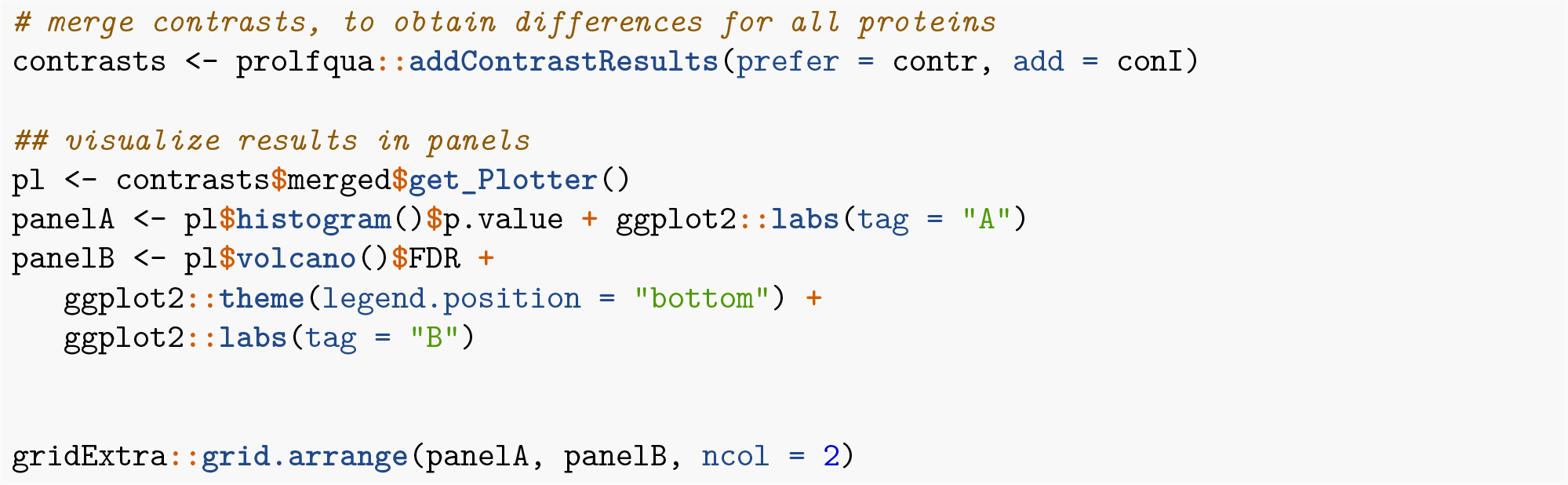

R linear model and linear mixed effect models allow modeling parallel designs, repeated measurements, factorial designs, and many more. Models in *prolfqua* are specified using R’s linear model and mixed model formula interface. Therefore, knowledge of the R regression model infrastructure (Faraway 2016; Venables and Ripley 2002) is advantageous when using our package. Furthermore, this glass box approach should make it easy to reimplement an analysis performed with *prolfqua* using base R or other programming languages by reading the analysis script. However, acknowledging the R formula interface’s complexity to non-statisticians and the popularity of *MSstats*, we provide the functionality to suggest a model formula from the sample annotation provided in tabular form similarly to *MSstats*.

Using the tidy table to model the data ensures interoperability with other proteomics-related packages that manage their data with tidy-tables, e.g., *protti* (Quast, Schuster, and Picotti 2022). The use of R6 classes, which encapsulate the configuration and the data, allows for writing very concise code (no function arguments needed). Auto-completion support in the Rstudio editor makes it easy for novices to explore *prolfqua*s functionality (Figure 6). To simplify integration of *prolfqua* with Bioconductor based workflows there is a method to convert the LFQData class into a *SummarizedExperiment*.

To ease the usage barriers of the R package to users not being familiar with statistics and R programming, we integrated some workflows into our data management platform b-fabric (Türker et al. 2010). This integration enables our users to select the input and basic settings in a graphical user interface. Then, as an output, the user receives a report including input files, the R markdown file, and R scripts necessary to replicate the analysis using their in-house R installation. The b-fabric system runs a computing infrastructure controlled by a local resource management system that supports cloud-bursting (Aleksiev et al. 2013). In this way, b-fabric helps scientists to meet requirements from funding agencies, journals, and academic institutions to publish data according to the FAIR (Findable, Accessible, Interoperable and Reusable) (Wilkinson et al. 2016) data principles.

### 3.2 Benchmarking modelling approaches

The Benchmark functionality of *prolfqua* includes receiver operator curves (ROC) and computes partial areas under those curves (pAUC) and other scores, e.g., the false discovery proportion FDP. We use those scores, i.e. the *pAUC* at 10 FDR and the FDP, to examine how well the methods implemented in *prolfqua* model quantitative mass spectrometric high throughput data and compare them with results produced by *MSstats* and *proDA*. Table 4 summarizes all modelling methods we evaluated.

**Table 3:** Prolfqua functions to fit various models.

| prolfqa functions | model |
| --- | --- |
| strategy_lm, Contrasts | linear modelling of peptide or protein abundances and group difference estimation |
| strategy_lmer, Contrasts | mixed effect modelling of peptide or protein abundances and group differences estimation |
| ContrastsSimpleImpute | group difference estimation when no observations in one of the groups |
| ContrastsROPECA | estimating group differences for proteins by summarizing peptide differences |
| ContrastsModerated | empirical Bayes variance shrinkage for group difference estimates (limma) |
| runSaint, ContrastsSaintExpress | protein interaction scoring (SAINTExpress) |
| strategy_proDA*, Contrasts_proDA* | adapter to the probabilistic dropout model implemented in proDA |

<sup>a</sup> in development

**Table 4:**
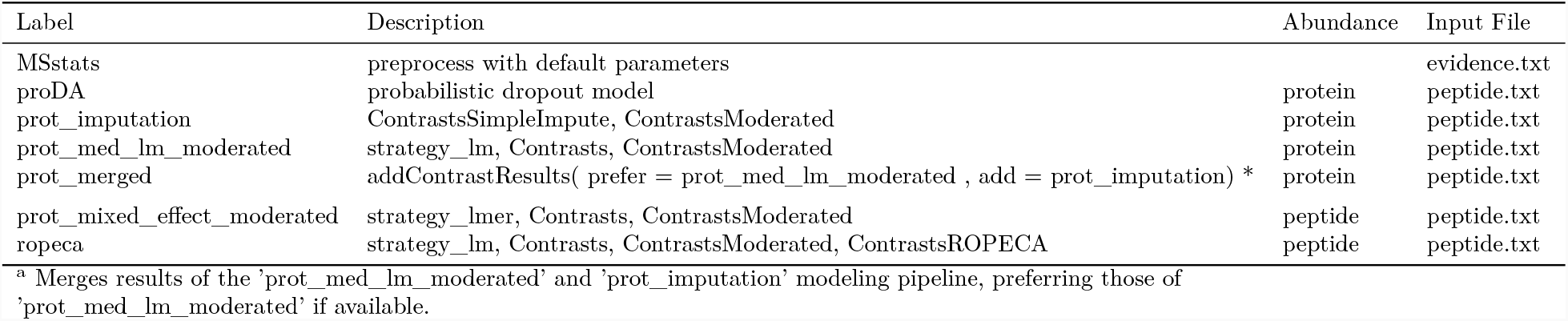
All benchmarked models. Description - prolfqua function names, Abudances - indicates if model is fitted to peptide or protein abundances, Input File - name of MaxQuant file used as input.

| Label | Description | Abundance | Input File |
| --- | --- | --- | --- |
| MSstats | preprocess with default parameters |  | evidence.txt |
| proDA | probabilistic dropout model | protein | peptide.txt |
| prot_imputation | ContrastsSimpleImpute, ContrastsModerated | protein | peptide.txt |
| prot_med_lm_moderated | strategy_lm, Contrasts, ContrastsModerated | protein | peptide.txt |
| prot_merged | addContrastResults( prefer = prot_med_lm_moderated , add = prot_imputation) * | protein | peptide.txt |
| prot_mixed_effect_moderated | strategy_lmer, Contrasts, ContrastsModerated | peptide | peptide.txt |
| ropeca | strategy_lm, Contrasts, ContrastsModerated, ContrastsROPECA | peptide | peptide.txt |

<sup>a</sup> Merges results of the 'prot\_med\_lm\_moderated' and 'prot\_imputation' modeling pipeline, preferring those of 'prot\_med\_lm\_moderated' if available.

When comparing differential expression analysis performance, a relevant parameter is the number of proteins for which a method estimated differences (see Figure 5 Panel A), which indicates how robust the procedure works in the presence of missing observations. For each protein, we tried to determine four differences (Δ = (1.20, 1.25, 1.30, 1.50)). Given 4046 proteins with at least two peptides, there are in total 16184 possible differences. However, some methods can not estimate all of them. The set of proteins with effect size estimates might differ for each method, biasing direct comparison of scores such as pAUC. For instance, we observe that *MSstats* estimates 16058 group differences while the mixed effect models estimates the fewest with 15940. Hence, to conclude that one method shows a better performance, it does not suffice if the pAUC is greater, but the number of proteins with differential expression results needs to be equal or larger.

**Figure 5:**
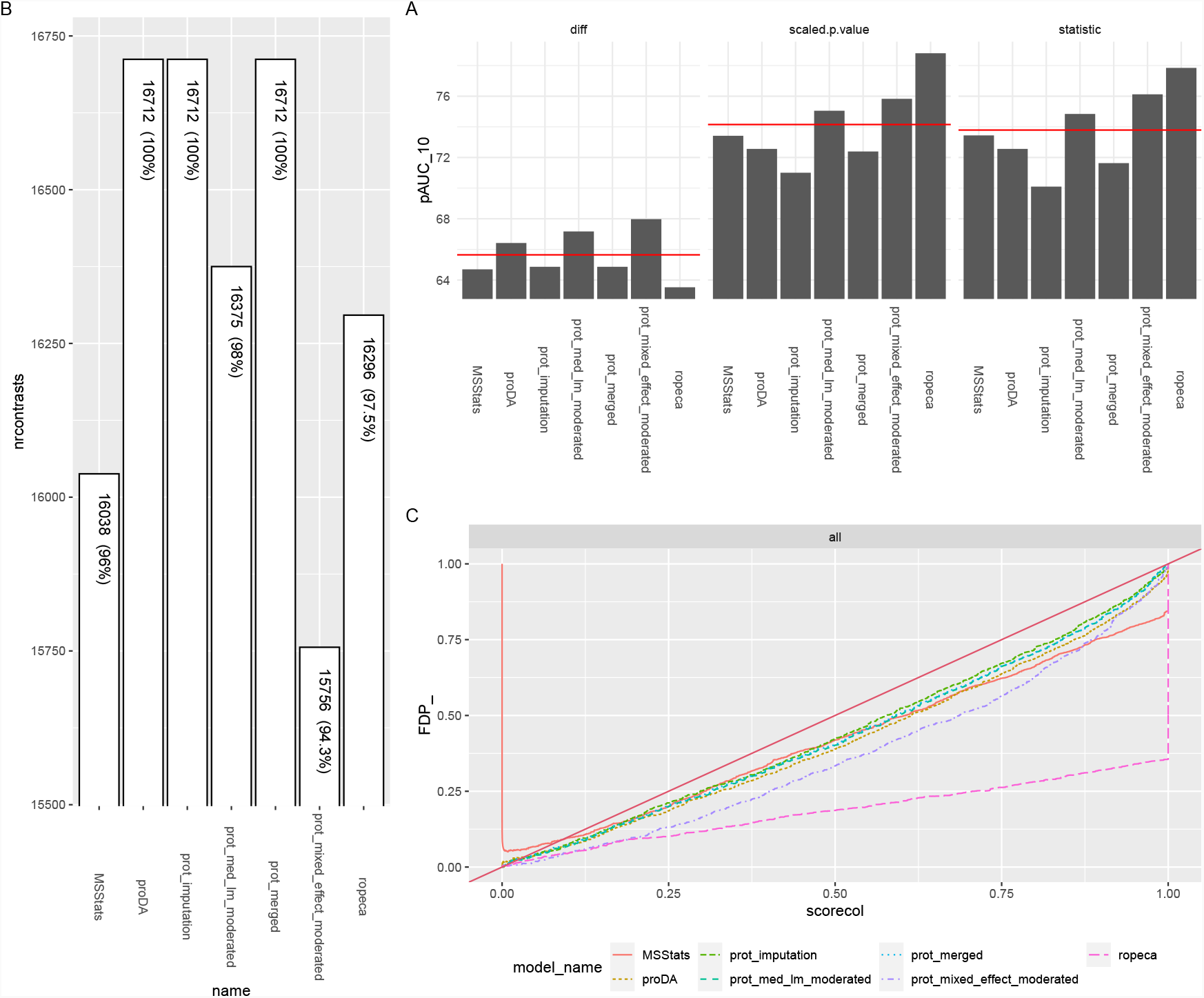
Panel A - Number of estimated contrasts for each modeling method (higher is better). Panel B - Partial area under the ROC curve at 10% FPR (*pAUC*_1_0) for all contrasts and three different statistics: the difference among groups, the scaled *p*-value (sign(diff) p.value) and the *t*-statistics (higher is better). The red line indicates the average area under the curve of all methods. Panel C - Plots the false discovery proportion (FDP) as a function of the FDR. Ideally, the FDR should be equal to the FDP. Therefore larger distances from the diagonal are worse.

Figure 5 Panel B shows how various estimates obtained from the models, i.e., the difference between groups, *t*-statistics, and scaled *p*-values allow identifying true positives (TP) given a false positive rate (FPR) of 10, by displaying the partial area under the ROC. Ordering proteins by the *t*-statistic or *p*-value leads to a higher *pAUC*_10_ than when ordering by the estimated difference among groups. Suppose an accurate estimate of the difference among groups is essential. In that case, the linear models fitted to protein intensities, calculated using Tukey’s median polish, perform better than those directly modeling peptide intensities, e.g., `ropeca` or `prot_mixed_effect_moderated` (see Table 4 Abundance column). We speculate that outliers at the peptide level do not affect the protein estimates when using Tukey’s median polish method, a non-parametric method to infer protein abundances. This hypothesis could be examined, by including other forms of protein intensity inferences implemented in `prolfqua,` e.g. `top-N` or `rlm,` into the benchmark.

There are only minor differences in the *pAUC*_10_ between the *t*-statistics or the scaled *p*-value (see Figure 5 Panel A). Interestingly, the *pAUC* increases when using *p*-values instead of the *t*-statistics for linear models, while it decreases for mixed effect models. The reason is an erroneous denominator degrees of freedom estimation for many proteins in the case of the mixed effect models.

We also benchmark if the *FDR* obtained from a model is an unbiased estimate of the false discovery proportion *FDP*. Figure 5 Panel C draws on the horizontal axis the FDR determined from the model, and on the vertical axis, the FDP obtained from the confusion matrix. Most lines are below the diagonal, indicating that the FDR estimates are modestly conservative for this benchmark dataset. In the case of *MSstats*, we observe a high proportion of false discoveries for small *FDR* values. In the case of *PECA*, the FDR estimates, obtained by Benjamini-Hochberg correcting the regulation probabilities, strongly overestimate the *FDP*.

Using a benchmark dataset whose ground truth is known (see Methods), we assessed the performance of different modeling approaches implemented in *prolfqua* (Tables 3 and 4), *MSstats* and *proDA*. Table 4 summarizes which methods we have evaluated, which MaxQuant results we used, and if the models are fitted to peptide or protein intensities. We make the R-markdown files to replicate the benchmarking available at *prolfquabenchamrk* and at BenchmarkingIonstarData.

Since only technical replicates are available for each dilution, essential sources of variation present in every experiment, such as the biochemical and biological, are not measured. Therefore, the dataset captures only the variance from the chromatography, electro-spray, and mass spectrometric measurement method. Thus, while we can extrapolate some of the results obtained to more realistic datasets, we need to be careful not to over-interpret the results. Specifically, the observed variances will be higher in more realistic experiments, and the power will be lower for the same sample sizes. Furthermore, the proportion of missing observations in real-life datasets might also be higher.

We can conclude that if we want to sort the proteins according to the likelihood of being differentially regulated to perform gene set enrichment analysis (Subramanian et al. 2005), the *t*-statistic is better suited than the fold-change estimate. Modeling the degrees of freedom when computing the *p*-values might improve the inference. However, this improvement is minuscule (see Figure 5 panel B). There is no such improvement for the mixed effect model, most likely because the degrees of freedom are erroneously estimated for many models. Furthermore, for the fixed effect linear model, the empirical Bayes variance shrinkage, as suggested by Smyth (2004), consistently improves the ranking of proteins compared with the unmoderated estimates (not shown) and fails to do so for mixed effect models.

Computing the statistics at the peptide level, e.g., the *t*-statistics or *p*-value, then summarizing these statistics using their median produces the highest *AUC* scores among all the tested models (see Figure 5 Panel A left). Furthermore, by using the Beta distribution to model the number of peptides observed, we can further improve the *pAUC* scores (see Figure 5 Panel A center). However, the properties of Beta-based probabilities are not well understood; for instance, the p-values are not uniformly distributed under the null hypothesis (not shown). Furthermore, the FDR estimates obtained when correcting for multiple testing with the Benjamini-Hochberg method are biased and overestimate the false discovery proportion (see Figure 5 Panel C). Therefore, we can not recommend this method if an unbiased estimate of FDR is essential, which is frequently the case. In addition, peptides are stronger affected by missing values, reducing the number of contrasts we could estimate for the dataset using this method (see Figure 5 Panel C).

The probabilistic dropout analysis implemented in the *proDA* produces inferences comparable to those of other methods (Figure 5 Panel A). Because of the robustness of the dropout model to missing observations we obtain difference estimates for all proteins and contrasts (Figure 5 Panel B). Moreover, the estimated differences are unbiased and show high diagnostic accuracy (Figure 5 Panel A). Furthermore, the performance of the scaled *p*-values or the *t*-statistics is comparable with that of the linear model with variance moderation `(prot_med_lm_moderated).` Therefore, we are planning to integrate the *proDA* package as an additional modeling option into *prolfqua* (see Figure 2).

The R-package *proDA* and *prolfqua* model the missing data directly, while MSstats imputes the data using an accelerated failure model. Despite imputation, *MSstats* does not estimate group differences for more proteins and does not achieve a higher *pAUC* score than *prolfqua*. Furthermore, Figure 5 Panel C shows that when using *MSstats* the proportion of false discoveries might be very high even when filtering for a low FDR.

We focused our benchmark on comparing the statistical modeling methods using three different scores, while we fixed the pre-processing steps. However, there are other equally or even more important parameters of a protein quantification pipeline (Fröhlich et al. 2022). One of them is the normalization of the intensities within the samples to remove systematic differences (Pursiheimo et al. 2015). The method used to infer protein intensities from peptide intensities is an additional important factor (Grossmann et al. 2010). For instance, the original *proDA* publication uses MaxLFQ (Cox et al. 2014) protein estimates. However, when using MaxLFQ intensities reported by MaxQuant, the *pAUC*_10_ is significantly lower (*pAUC*_10_(t-statistics) = 66) compared with results obtained when protein abundances are estimated from peptide abundances using Tukey’s median polish (*pAUC*_10_(t-statistics) = 73). Last but not least, the software (Cox and Mann 2008; Yu et al. 2020), identifying proteins and generating the quantification values can also significantly contribute to the performance of the entire pipeline, altering the number of identified proteins and the sensitivity and specificity of the differential expression analysis.

## 4 Conclusion

*prolfqua* allows for considerable flexibility to model quantitative proteomics experiments. Various types of models are available (see Figure 2 and Table 3), and the contrast specification is explicit and consistent for all models. The modular design of *prolfqua*, allows for adding new features, e.g., generalized linear models (*glm*’s) to model the presence or absence information of a protein, or robust linear models (*rlm*’s), in the future. R’s formula interface for linear models is flexible, widely used, and well documented (Faraway 2016). We use the formula interface to specify the models, making it easy to reproduce an analysis performed with *prolfqua* in other statistical methods programming languages. However, the developed framework is flexible enough to in the future integrate other modeling methods, e.g., the probabilistic dropout model (Ahlmann-Eltze and Anders 2020) or accurate variance estimation (Zhu et al. 2020). Hence, *prolfqua* enables you to call various methods and makes selecting the best differential expression analysis algorithm for your problem easy.

When comparing statistical modeling methods for the differential expression analysis, we assessed performance measures such as the number of estimated contrasts, the *pAUC*, and if the FDR is an unbiased estimate of the FDR. It is relevant that an analysis pipeline shows good performance in all these categories. Leveraging these computer experiments, we can provide the following advice: i) estimate protein abundances from peptide abundances using a robust or nonparametric regression method; ii) use linear models because they show good performance in all categories; iii) if the measurements are correlated, as for technical replicates, mixed effect models might work if the sample sizes are large; if not, aggregate the replicates and fit a linear model instead; iv) if you use fixed-effect linear models, apply variance moderation to improve the *t*-statistics and *p*-value estimates; v) If you want to sort your protein lists to perform gene set enrichment analysis, use the *t*-statistic instead of the difference; vi) do not impute missing observation but statistically model missingness to estimate parameters, i.e., group differences. Finally, the differential expression analysis result obtained with prolfqua are comparable to or better than when using other differential expression analysis tools.

In summary, *prolfqua* is an easy-to-use and feature-rich R package to analyze quantitative mass spectrometric data with simple or complex experimental designs. It also can generate conclusive reports and to benchmark MS software and statistical methods. Furthermore, with minimal adaptations, this R package can analyze different quantitative proteomics data (e.g., labeling-based TMT-, PRM-DIA-data). We provide documentation, in vignette format, at the website https://github.com/fgcz/prolfqua/. This document was created using Rmarkdown. All the code needed to replicate the document or the benchmark results is available at: https://github.com/wolski/prolfquabenchmark.

## Acknowledgements

The authors thank the technology platform fund (TPF) of the University of Zurich and all FGCZ proteomics colleagues for fruitful discussions.

## Abbreviations

Abbreviations: Explaination
AUC: Area Under the Curve
CDF: Cumulative Distribution Function
ESI-MS: Electro-Spray-Ionization Mass Spectrometry
*FDP*: False Discovery Proportion
*FDR*: False Discovery Rate
LC: Liquid Chromatography
LC-MS: Liquid Chromatography followed by Mass Spectrometry
LOD: Limit Of Detection
MAR: Missing At Random
MCAR: Missing Completely At Random MS mass spectrometry
OO: Object-Oriented
UML: Unified Modeling Language

## Appendix

### Creating a prolfqua configuration

The following code demonstrates how we use *prolfqua* to analyze protein intensities reported in the MSFragger `combined_protein.tsv` file. First, we create a tidy table containing the protein abundances by reading the `combined_protein.tsv` file using `tidy_MSFragger_combined_protein`. Then, we read the sample annotation from the file annotation.xlsx file. Next, we create an `AnalysisTableAnnotation` R6 object. Bottom-up proteomics data is hierarchical, i.e., a protein has peptides, peptides might be modified, etc. Therefore, the `AnalysisTableAnnotation` has a `hierarchy` field storing a list with an entry for each hierarchy level. Since combined_portein.tsv only holds protein level data, the hierarchy list has one element, and we use it to specify which column contains the protein identifiers. We also need to define which column contains the protein abundances we want to use for the data analysis. Finally, we have to specify which columns contain the explanatory variables of the analysis. The `AnalysisTableAnnotation` has the field `factors`, a list with as many entries as explanatory variables. Here we include two explanatory variables, the dilution, specified in the column ‘sample’, and ‘run’ stored in the column ‘run_ID’, representing the order of the measurement.

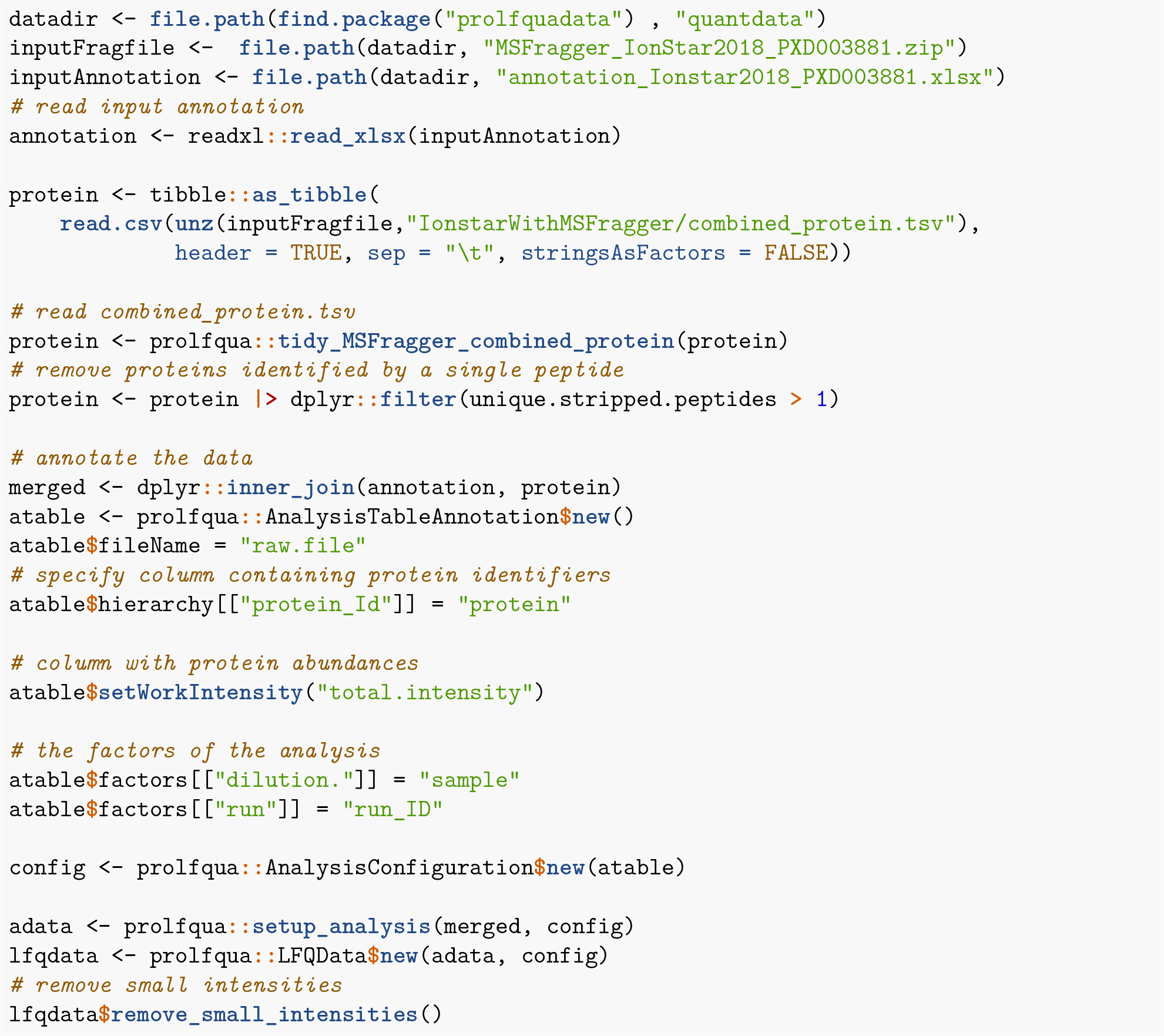

### Miscellaneous

**Figure 6:**
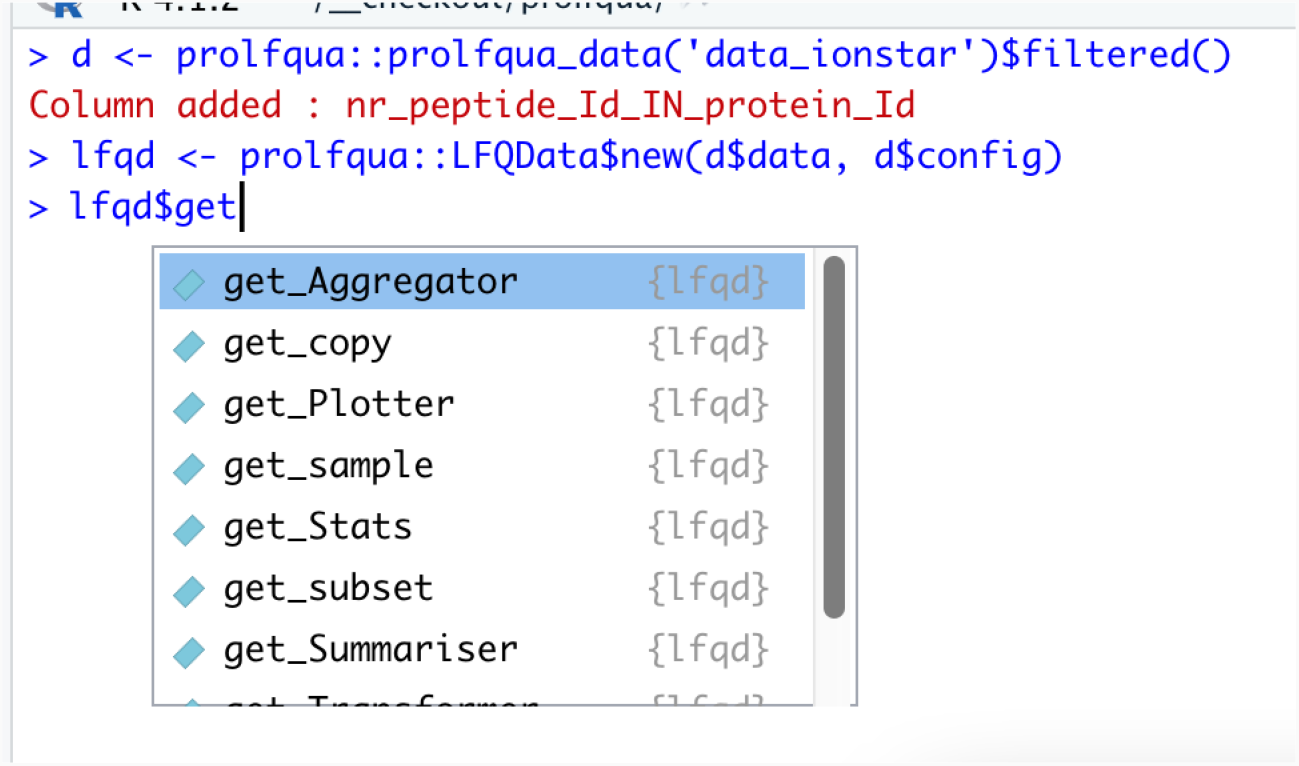
The screenshot displays the command-line completion (tab completion) of RStudio on the prolfqua::LFQData R6 object. In the example, it shows the getter methods of the object.

**Figure 7:**
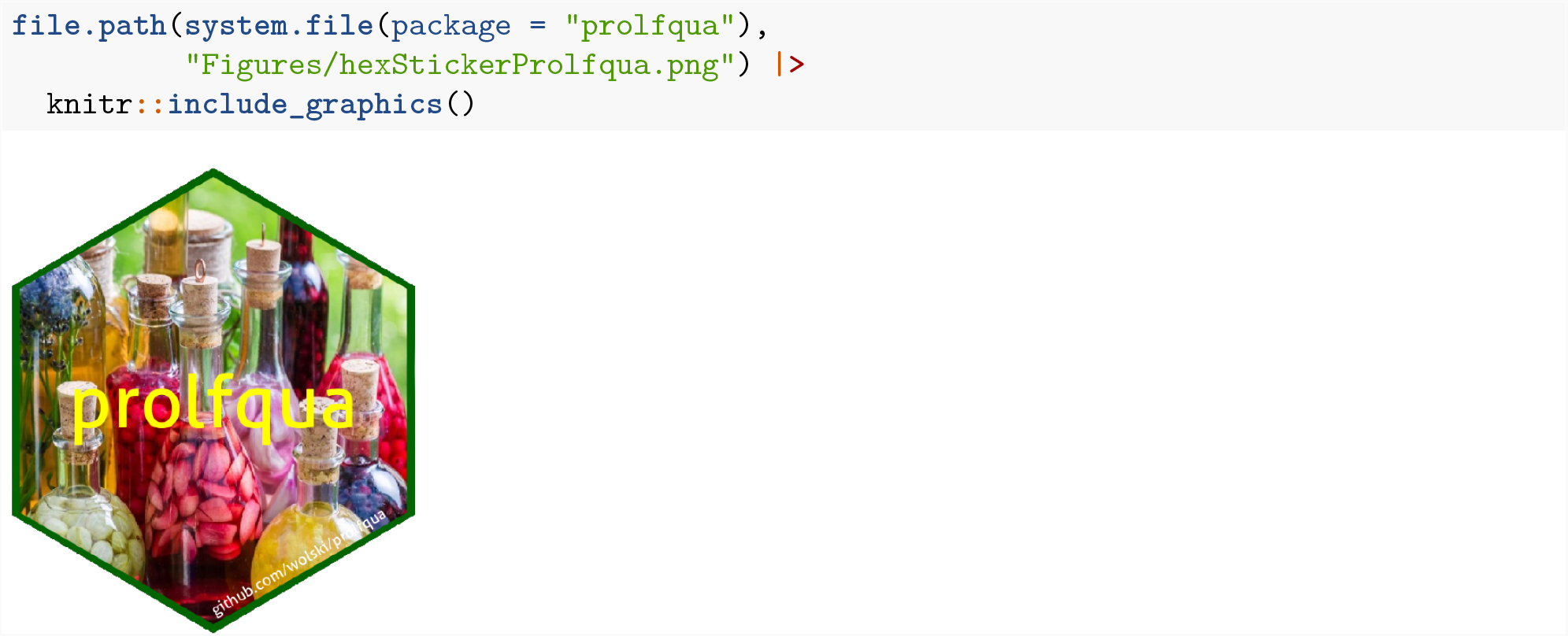
Sticker maintainer: Witold E. Wolski; License: Creative Commons Attribution CC-BY. Feel free to share and adapt, but don’t forget to credit the author.

### Session information

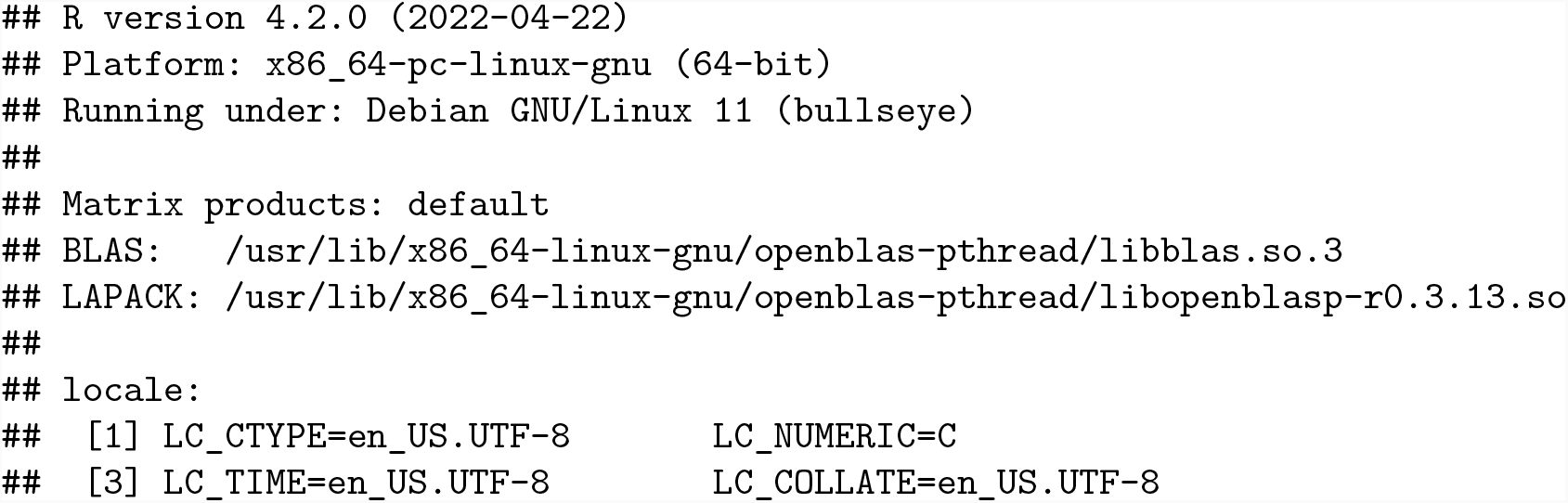
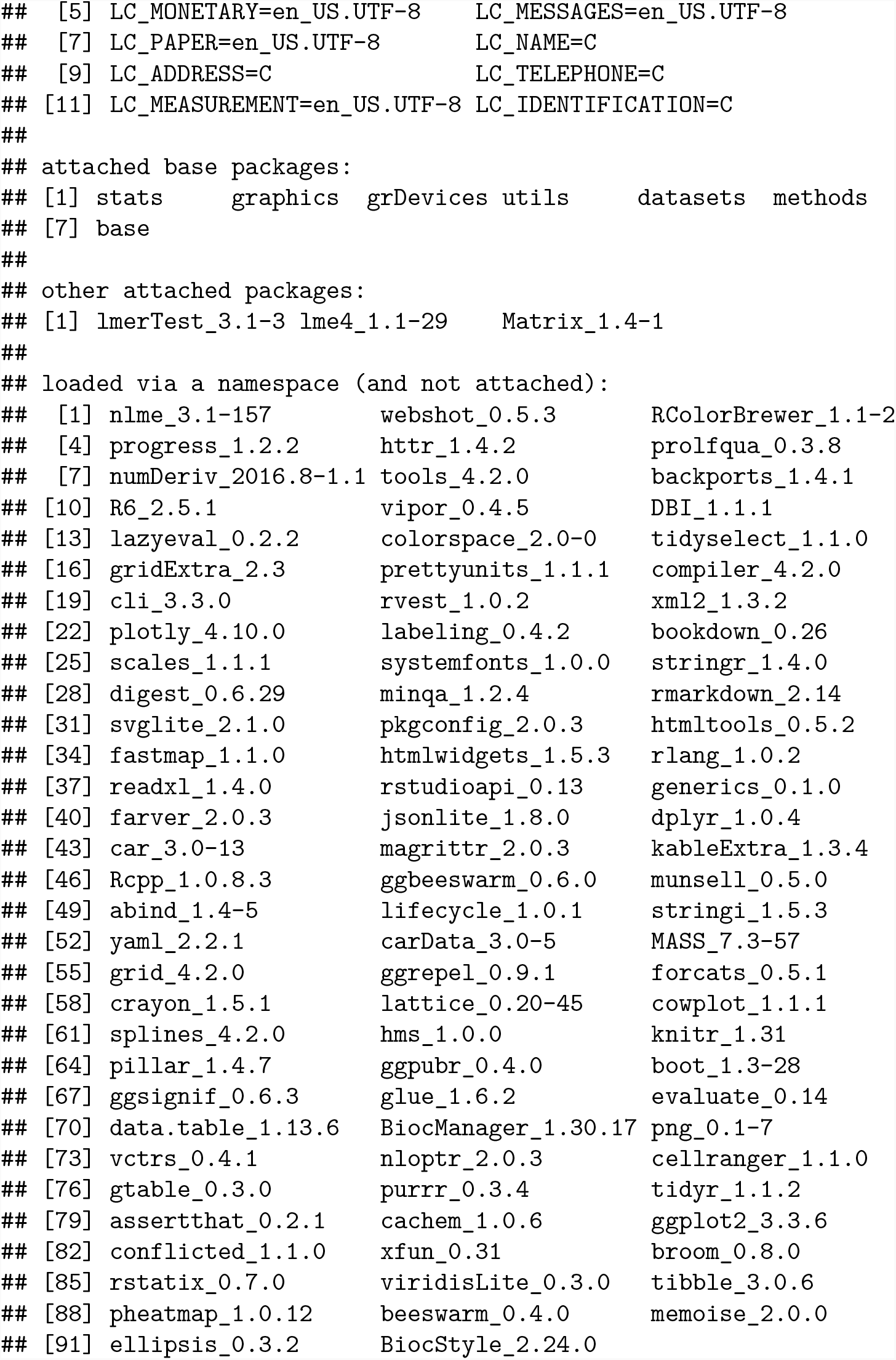

